# The genomic false shuffle: epigenetic maintenance of topological domains in the rearranged gibbon genome

**DOI:** 10.1101/238360

**Authors:** Nathan H. Lazar, Kimberly A. Nevonen, Brendan O’Connell, Richard E. Green, Thomas J. Meyer, Mariam Okhovat, Lucia Carbone

## Abstract

The relationship between evolutionary genome remodeling and the three-dimensional structure of the genome remain largely unexplored. Here we use the heavily rearranged gibbon genome to examine how evolutionary chromosomal rearrangements impact genome-wide chromatin interactions, topologically associating domains (TADs), and their epigenetic landscape. We use high-resolution maps of gibbon-human breaks of synteny (BOS), apply Hi-C in gibbon, measure an array of epigenetic features, and perform cross-species comparisons. We find that gibbon rearrangements occur at TAD boundaries, independent of the parameters used to identify TADs. This overlap is supported by a remarkable genetic and epigenetic similarity between BOS and TAD boundaries, namely presence of CpG islands and SINE elements, and enrichment in CTCF and H3K4me3 binding. Cross-species comparisons reveal that regions orthologous to BOS also correspond with boundaries of large (400-600kb) TADs in human and other mammalian species. The co-localization of rearrangement breakpoints and TAD boundaries may be due to higher chromatin fragility at these locations and/or increased selective pressure against rearrangements that disrupt TAD integrity. We also examine the small portion of BOS that did not overlap with TAD boundaries and gave rise to novel TADs in the gibbon genome. We postulate that these new TADs generally lack deleterious consequences. Lastly, we show that limited epigenetic homogenization occurs across breakpoints, irrespective of their time of occurrence in the gibbon lineage. Overall, our findings demonstrate remarkable conservation of chromatin interactions and epigenetic landscape in gibbons, in spite of extensive genomic shuffling.

## Introduction

The spatial organization of a genome and its chromatin interactions play a crucial role in mediating vital cellular functions, such as gene regulation (Lieberman-Aiden et al., 2009) and DNA replication (Pope et al., 2014). Intra-genomic interactions organize chromosomes into functional compartments called topologically associating domains (TADs). Within TADs, nearby loci (i.e. enhancers and genes) interact more frequently with each other than with regions located elsewhere in the genome (Dixon et al., 2012). Genes located in the same TAD are often co-regulated, co-expressed, and display correlation in epigenetic marks of chromatin activity (Nora et al., 2012). Genes and regulatory elements in neighboring TADs are epigenetically and functionally insulated by TAD boundaries. The zinc finger CCCTC-binding factor (CTCF) is thought to play a role in TAD boundary formation and mediation of long-range chromatin interactions; however other factors, including the level of transcriptional activity and presence of retrotransposons, may also contribute to boundary formation (Dixon et al., 2012).

TADs appear to be largely conserved in their structure and organization across tissues and species, as the majority (53.8%) of TADs have been found to be evolutionarily conserved between human and mouse embryonic stem cells (Dixon et al., 2012). Moreover, visual pairwise comparisons of genome-wide chromatin conformation capture (Hi-C) and CTCF binding data between mouse, rhesus, dog, and rabbit showed that overall TAD structure is maintained in syntenic regions (Vietri Rudan et al., 2015). Interestingly, conservation of TAD organization and structure appears to strongly relate to TAD size: large TADs (>1Mb) show higher conservation across cell types and species, while sub-TADs (100kb-1Mb) are more variable and contribute to cell- or species-specific gene regulation (Dixon et al., 2012; Phillips-Cremins et al., 2013; Rao et al., 2014). Despite recent advancement in the understanding of TAD structure and function, the mechanisms behind evolutionary TAD conservation have yet to be fully explored.

Chromosomal rearrangements (deletions, duplications, translocations, and inversions) are large-scale events that can drastically influence TAD integrity and organization during species evolution. Until now, studies that have examined the consequences of chromosomal rearrangements on TADs have primarily focused on pathological structural variations and therefore have been limited to discrete genomic regions. Using genetic engineering in patient cell lines and mouse models, these studies show that rearrangements that remove or misplace TAD boundaries can initiate ectopic interactions between genes and regulatory elements of neighboring TADs, leading to aberrant phenotypes and pathology (e.g. limb malformations, leukemia) (Dixon et al., 2017; Franke et al., 2016; Groschel et al., 2014; Hnisz et al., 2016; Lupianez et al., 2015). However, there is a paucity of information on the evolution of genome topology including how long-distance DNA interactions and the epigenetic landscape of TADs change following evolutionary rearrangements. With the exception of one study (Vietri Rudan et al., 2015) previous comparative work mainly focused on broad comparisons of TADs between human and mouse, and leaves out comparisons between more closely related species (Dixon et al., 2012; Rao et al., 2014). Moreover, these studies lack a detailed and quantitative assessment of the relationship between synteny breakpoints and the epigenetic structure of TADs.

Effects of chromosomal rearrangements on TAD organization and gene regulation during genome evolution will depend on the location of rearrangement breakpoints. Rearrangements occurring at domain borders will maintain TAD integrity and preserve gene regulation within these units (Figure 1). However, these events might still modify the physical positioning of loci within the nucleus and perturb long-distance *cis* and *trans* interactions between nuclear territories (Harewood and Fraser, 2014). Rearrangements that disrupt TADs by breaking the DNA within boundaries might generate new functional interactions and even new TADs (Figure 1). These rearrangements can cause disease (Redin et al., 2017; Zepeda-Mendoza et al., 2017) making them likely to be eliminated by purifying selection because of their negative effect on fitness. However, a small portion of new TADs might contribute to new traits, giving rise to evolutionary novelties favored by selection.

**Figure 1.**
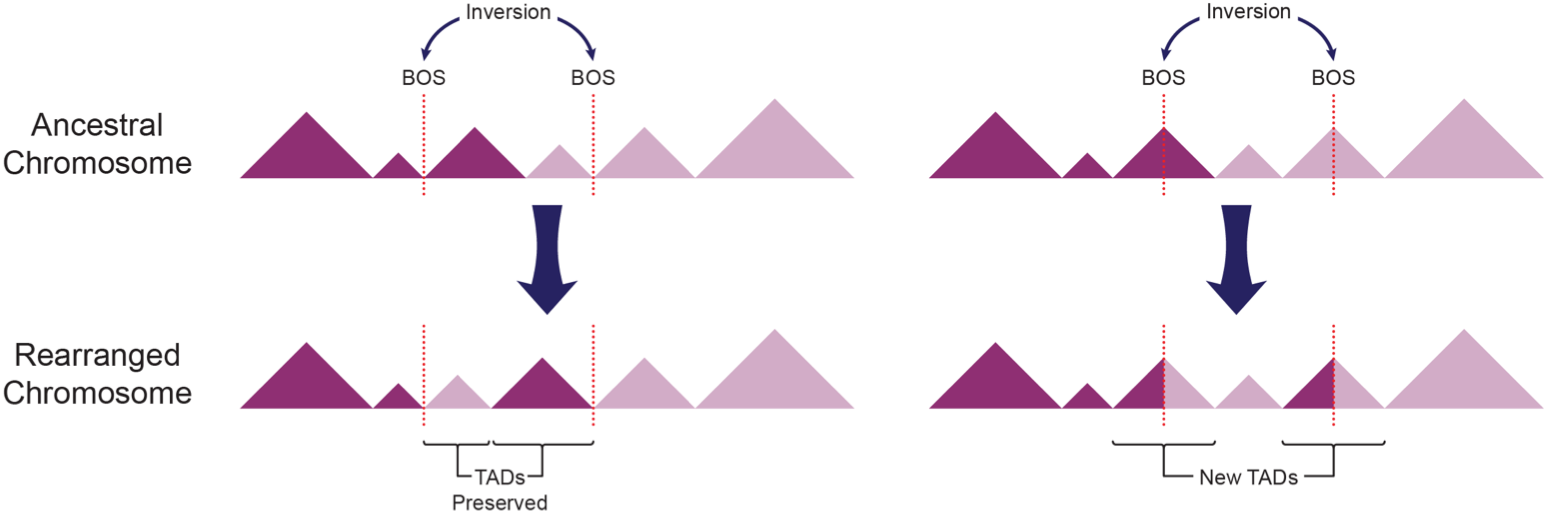
Relative position of breaks of synteny differentially affects TAD integrity. Alternative consequences of hypothetical ancestral inversions (top) are demonstrated. TADs are represented with triangles and positions of breaks of synteny (BOS) are depicted with dotted lines. A) BOS occurring at TAD boundaries in the ancestral genome, rearrange TADs as intact modules. B) BOS within TAD bodies disrupt ancestral TADs and may give rise to new TADs in the gibbon genome.

Gibbons are particularly appealing experimental models for studying the relationship between TADs and evolutionary chromosomal rearrangements, as this group has experienced an unusually high number of chromosomal rearrangements since their relatively recent divergence from humans (~17 million years ago). In addition, a high quality, well-annotated genome is available for the northern white-cheeked gibbon (*Nomascus leucogenys*, NLE) (Carbone et al., 2014) along with a high-resolution identification of evolutionary breaks of synteny with the human genome, validated by both DNA sequencing and molecular cytogenetics (Capozzi et al., 2012; Carbone et al., 2009a; Carbone et al., 2006; Girirajan et al., 2009; Roberto et al., 2007). Finally, the high genetic identity between gibbon and human (~96%) enables the use of many genetic, epigenetic, and computational tools designed for use in human studies and facilitates the comparison between the two species.

In this study we use the gibbon genome to examine the evolutionary consequences of heavy genome reshuffling on long distance DNA interactions and the integrity of TADs and their boundaries. We observe and quantify a striking co-localization of gibbon-human breaks of synteny (BOS) with TAD boundaries, independent of parameters used to identify TADs. This observation is further supported by the genomic and epigenetic signatures found at gibbon breakpoints including the presence of CTCF binding, H3K4me3, enrichment of SINE elements, and CpG islands. Our findings suggest that the extensive chromosomal remodeling observed in the gibbon happened within the constraints of genome topology. Thus, although the gibbon genome is structurally shuffled, its regulatory units remained largely intact and conserved with respect to the hominoid ancestor.

## Results

### Gibbon Hi-C interaction map highlights structural variations and recent genomic rearrangements

Whole-genome maps of chromatin interactions can be used to identify and study non-reference structural variations in a genome (Dixon, 2017; Harewood et al., 2017). In order to characterize the relationship between gibbon chromosomal rearrangements and TAD organization, we performed genome-wide chromatin conformation capture sequencing (Hi-C) using a lymphoblastoid cell line (LCL) that we established from a male northern white-cheeked gibbon (*Nomascus leucogenys*, Vok, #NLL600) belonging to the same species used to generate the gibbon genome reference (Carbone et al., 2014). Our Hi-C data, exhibited evidence for a translocation between gibbon chromosomes 1 and 22, which is a known polymorphic rearrangement in the genus *Nomascus* (Koehler et al., 1995) (Figure 2A). The individual used to generate the Hi-C data appears to carry the ancestral forms, NLE1a (corresponding to human chromosomes 9, 6 and 2) and NLE22a (corresponding to human chromosome 14), whereas the reference genome contains sequences for the derivative NLE1b (corresponding to human chromosomes 9, 6 and 14) and NLE22b (corresponding to human chromosomes 14, 6 and 2). In our Hi-C map, this non-reference translocation appears as strong inter-chromosomal interactions in conjunction with absence of intra-chromosomal interactions. We confirmed this translocation by fluorescent *in situ* hybridization (FISH) using whole-chromosome paints for NLE1b and NLE22b on Vok chromosome spreads (Figure 2B **and Supplemental Fig. S1**).

**Figure 2.**
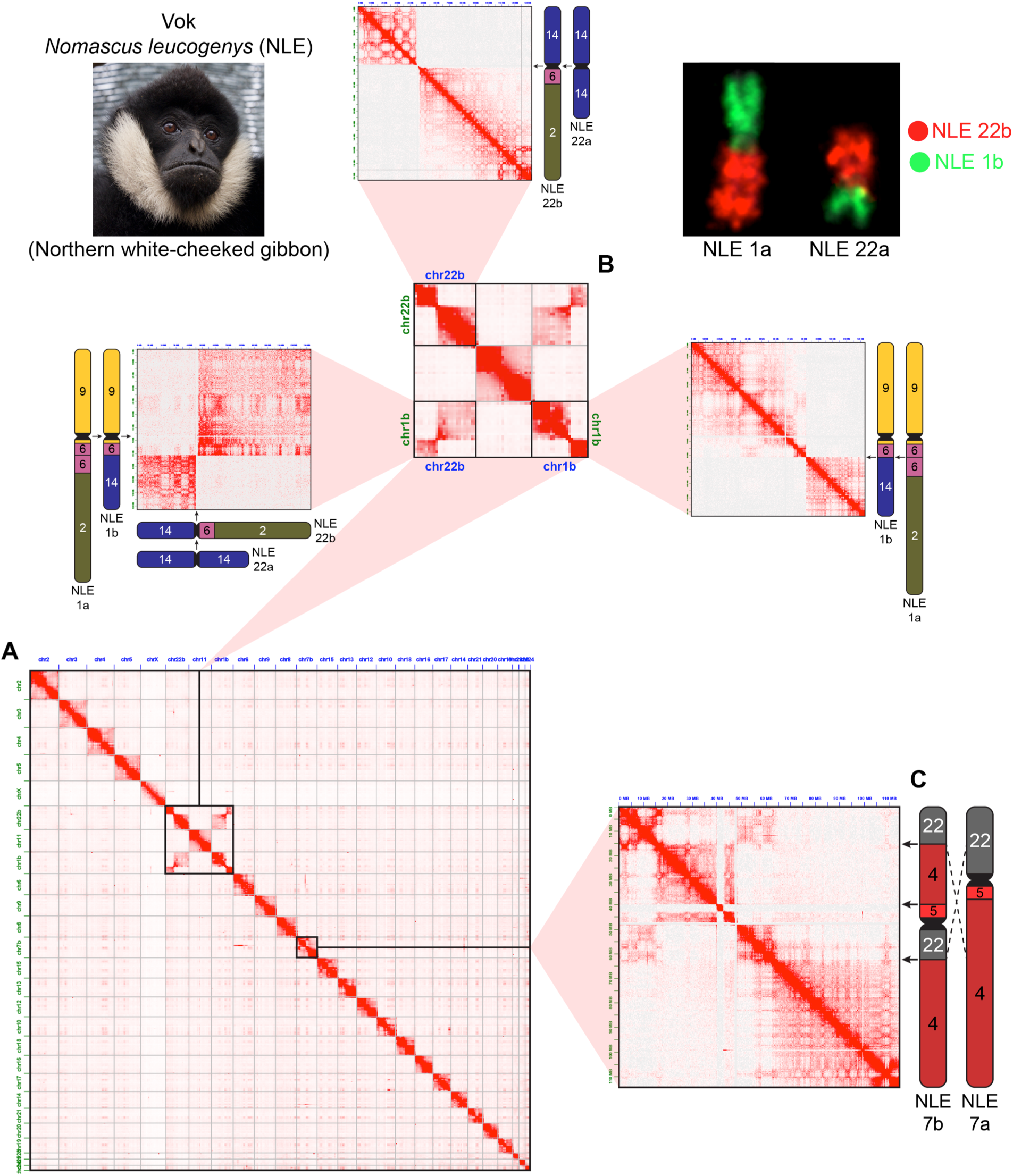
Gibbon Hi-C map highlights non-reference chromosomal rearrangements. A) Whole genome Hi-C interaction matrix for the gibbon named Vok (photo shown in the upper left corner), aligned to the reference gibbon genome (Nleu3.0). B) Zoomed image of the reciprocal translocation between chr1 and chr22, indicated by a lack of intra-chromosomal interactions in the regions mobilized by the rearrangement, and stronger-than-expected inter-chromosomal interactions. Vok carries the ancestral forms (NLE1a/22a) while the reference genome carries derivative forms (NLE1b/22b). FISH with chromosome paints for NLE1b (green) and NLE22b (red) on Vok chromosomes is shown on the upper right corner. C) Hi-C matrix for NLE7b displays examples of ‘ghost interactions’ between regions corresponding to human chromosome 22, separated by a recent pericentromeric inversion in the ancestral NLE7a. Gibbon chromosomes are labeled outside of chromosome ideograms and corresponding human chromosomes are indicated within ideograms.

Other interaction signals suggestive of non-reference translocations were also observed. For instance, we identified strong interactions between a 2Mb “strip” on NLE7b (chr7b: 39,750,000-42,500,000) and the entire NLE6 **(Supplemental Fig. S2A)**, while the same strip barely interacted with other regions in *cis*. Similar patterns of unusual inter-chromosomal interaction were found in Hi-C data of rhesus, rabbit and dog **(Supplemental Fig. S2B)**. It should be noted however, that these putative translocations were not experimentally validated and may simply reflect errors in the genome assemblies.

Interestingly, we observed that genomic regions that were syntenic in the ancestral chromosome sometimes maintained their interaction even after loss of synteny occurred in gibbon due to rearrangement. These “ghost interactions” were mainly detected in recent evolutionary rearrangements with the most prominent case being present in gibbon chromosome NLE7b, whose current structure resulted from the most recent (~2 mya) rearrangement in the genus *Nomascus*. This rearrangement is a pericentromeric inversion of the ancestral form (NLE7a) (Figure 2C) which is only present in the northern white-cheeked gibbon (Carbone et al., 2009b). Our Hi-C interaction data showed that regions of NLE7b corresponding to human chromosome 22 still interact strongly with one another despite being separated on different chromosome arms in the new genomic arrangement (Figure 2C). We also observed a few cases of weaker, yet visually evident, inter-chromosomal ghost interactions. Namely, we observed stronger-than-background interactions between portions of NLE14 and NLE19 homologous to human chromosome 17. Moreover, some regions that were broken up by recent evolutionary rearrangements, still display similar patterns of interaction with other chromosomes in *trans*. For example, the chromosomal segments in NLE7 that are homologous to human chromosome 22 all display strong inter-chromosomal interactions with NLE14 and NLE6, while overall, the rest of NLE7 exhibits weaker interaction with these chromosomes **(Supplemental Fig. S2A)**.

### Gibbon-human breaks of synteny co-localize with gibbon TAD boundaries

Here we define a break of synteny (BOS) as a locus of the gibbon genome whose 5’ and 3’ ends are homologous to different, non-syntenic segments of the human genome **(Supplemental Table S1)**. Each gibbon BOS therefore corresponds to two distinct regions in the human genome, either located on the same chromosome (in the case of inversions) or on two different human chromosomes (in the case of translocations, fissions, or fusions). The human genome was selected to represent the ancestral hominoid genome because of its unmatched quality amongst primate genome assemblies. The rhesus macaque genome was used to discriminate human- and great ape-specific from gibbon-specific rearrangements (Carbone et al., 2009a). We characterized a total of 67 gibbon BOS. Among these, 33 were identified at single base-pair resolution, while the remaining 34 BOS contained insertions of repetitive elements of variable sizes that lack synteny with the human genome, and were therefore defined as “intervals”. All of these BOS regions have been previously validated by FISH and Sanger sequencing (Carbone et al., 2009a; Girirajan et al., 2009) and represent a highly curated set that excludes ambiguous mapping or genomic regions with assembly issues.

Alignment of gibbon Hi-C contact maps against the UCSC browser gibbon-human chain, allowed us to visually inspect the correspondence between BOS and genomic interactions. We observed an overall reduction of chromatin interactions across gibbon BOS (Figure 3A). To examine whether this was a general pattern, we averaged Hi-C data surrounding all BOS (+/−2.5Mb) into a single two-dimensional contact map using Homer (Heinz et al., 2010) (Figure 3B). We found a clear reduction in interactions between the two sides of the overlaid BOS, a pattern characteristic of TAD boundaries. Repeating this process with randomly generated regions with the same chromosome distribution and size of the gibbon BOS, did not produce a similar signal(Figure 3B).

**Figure 3.**
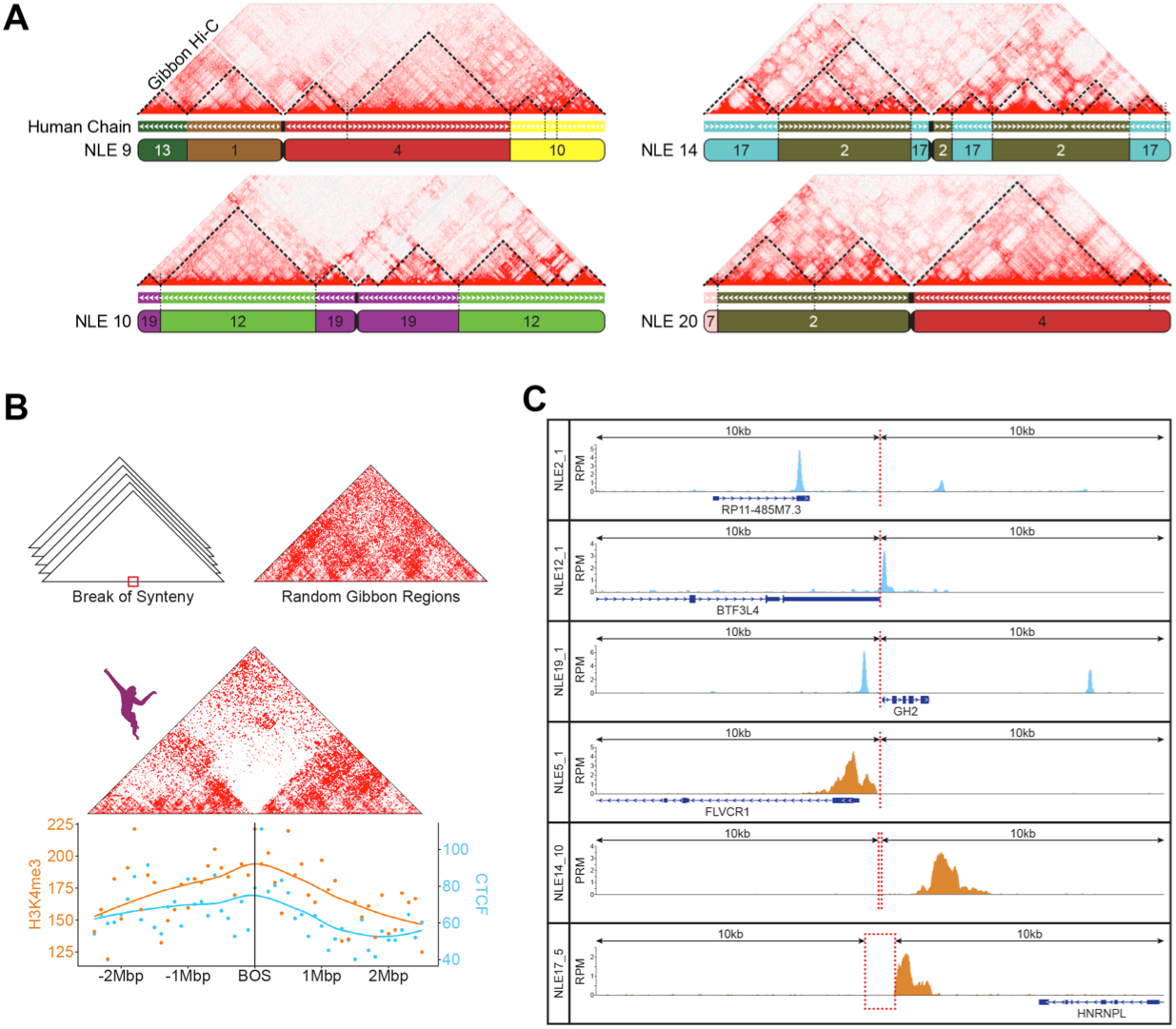
Gibbon-human breaks of synteny display epigenetic signatures of TAD boundaries. A) Hi-C matrix for four representative gibbon chromosomes (NLE9, 10, 14 and 20) aligned with the gibbon-human chain from the UCSC browser shows overlap between putative TAD boundaries and gibbon-human BOS. Corresponding human chromosomes are color coded and labeled within each gibbon chromosome ideogram. B) Top, two-dimensional histograms show juxtaposition of the Hi-C signal from regions flanking the BOS (+/−2.5Mbp) in gibbon and flanking random genomic regions (top right corner). Chromatin contacts are depleted across BOS, but not random regions. Bottom, CTCF and H3K4me3 ChIP-seq peak counts with smoothed Loess curves in 100kb bins across the BOS (+/− 2.5Mb) show enrichment of these epigenetic marks at BOS. C) Examples CTCF (blue) and H3K4me3 peaks (orange) in a 20Kb window around BOS, RPM= Reads Per Million [i.e. (reads aligned)/(million read sequenced)].

TAD boundaries are known to be occupied by CTCF as well as associated with epigenetic marks of active transcription (i.e. H3K4me3) (Dixon et al., 2012). Therefore, if gibbon BOS co-localize with TAD boundaries, we expect to observe the same epigenetic marks in these regions. To this end, we analyzed newly generated H3K4me3 ChIP-seq data for Vok and the gibbon used for the reference genome (Asia) and used previously published gibbon CTCF ChIP-seq data (Carbone et al., 2014). Although more significant for CTCF, we found significant enrichment for both marks within 20kb of gibbon BOS (two-sided permutation p-values, CTCF: <0.001, H3K4me3: 0.034) (Figure 3B-C). We also found that CpG density in the 20kb surrounding the gibbon BOS is significantly higher than the rest of the genome (two-sided permutation p-value: <0.001) **(Supplementary Material and Supplemental Fig. S3-S6)** and that BOS are significantly enriched in CpG islands, CpG shores, and SINE elements **(Supplemental Fig. S6)**. These genomic features have been described as characteristic of TAD boundaries in human and mouse (Dixon et al., 2012; Sun et al., 2017).

Next, we used our Hi-C data to computationally predict the position of gibbon TAD boundaries and examine if they significantly overlap with BOS. Since size and position of predicted TAD boundaries can vary depending on the algorithm and parameters used, we predicted TADs with 140 different parameter combinations (7 matrix resolutions, 5 cutoffs, and 4 window sizes; Supplemental Material) using the directionality index method (Dixon et al., 2012) and TADtool (Kruse et al., 2016). As expected, we observed that parameter choices greatly impacted the location and size of TAD boundaries **(Supplemental Fig. S7)**. In order to quantify the overlap of TAD boundaries with BOS in an unbiased way, for each set of parameters, we performed permutation analyses comparing the number of BOS-TAD boundary overlaps to the number of overlaps of TAD boundaries with 10,000 sets of random regions that had the same size and chromosomal distribution as the BOS regions. We found a significant correspondence (permutation p-values<0.001) between gibbon BOS and TAD boundaries for all resolution values, except the two extremes (10kb and 1Mb) and it was independent of the window size and cutoff **(Supplemental Fig. S8)**.

### Regions orthologous to gibbon BOS co-localize with boundaries of larger TADs in other mammalian species

To better understand the evolutionary relationship between gibbon TAD boundaries and chromosomal rearrangements, we examined genomic regions orthologous to gibbon BOS **(Supplemental Table S1)** and TAD structures in five other species (human, rhesus, mouse, dog, and rabbit) using publicly available, species-specific Hi-C data (Grubert et al., 2015; Vietri Rudan et al., 2015). First, we normalized and overlaid the Hi-C data from these species to examine DNA interactions +/− 2.5Mbp around loci orthologous to gibbon BOS (Figure 4A). Since each side of a gibbon BOS maps to a disjoint region, there were roughly twice as many regions composing each map, with some regions missing, because they failed to lift over from the gibbon genome. Interestingly, all species showed a distinct reduction in contacts across the sites orthologous to gibbon BOS, suggesting that loci where rearrangements occurred in gibbon are more likely to be TAD boundaries in other species (Figure 4A).

**Figure 4.**
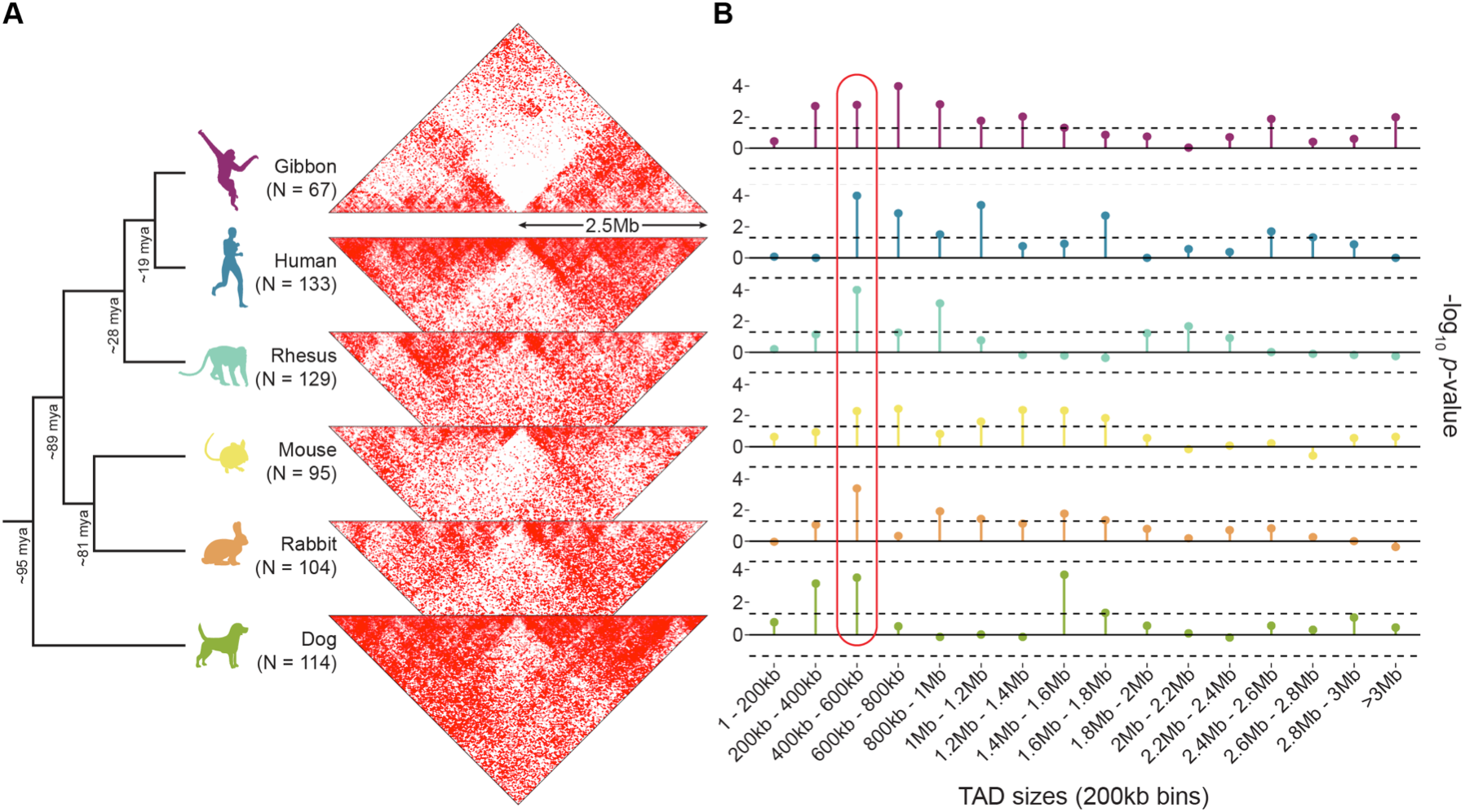
Evolutionary context of the overlap between TAD boundaries and BOS. A) The two-dimensional gibbon Hi-C histogram (Fig. 3B) is compared with Hi-C histograms for five other mammalian species at loci orthologous to the gibbon BOS (+/− 2.5Mbp). Decreased contact density across these loci in non-gibbon species suggests that breakpoint regions in gibbon are more likely to be TAD boundaries in other species. B) Lollipop plots show −log 10 p-values from permutation analyses testing the overlap between gibbon BOS and TADs binned by size. This cross-species comparison points to consistently significant overlap of BOS with boundaries of 400-600kb TADs (circled in red). Dotted line marks p=0.05 significance threshold (no multiple-test correction).

To predict TAD boundaries in all five species we used TADtool (Kruse et al., 2016) with the 140 different parameter combinations previously used in gibbons (data not shown). We detected a significant overlap between TAD boundaries and regions orthologous to gibbon BOS across all species for the majority of parameters combinations **(Supplemental Fig. S8)**. Expectedly, the association was more consistent in the gibbon genome, which is likely due to independent species-specific remodeling events in each lineage. The weakest overlap was observed in dog, possibly due to the highly rearranged nature of the dog genome compared to other mammals (Webber and Ponting, 2005).

Several studies indicate that larger TADs (>1Mb) show more conservation across species and tissues, whereas smaller subdomains (100kb-1Mb) appear more dynamic and change during development (Dixon et al., 2012; Nora et al., 2012; Phillips-Cremins et al., 2013). In order to facilitate comparison across species and establish the relationship between BOS and boundaries based on TAD size, we repeated our permutation analyses after grouping TADs by size, rather than parameter combinations **(Supplemental methods)**. First, TADs were grouped based on size (1bp-50kb, 50-100kb, 100-250kb, 250-500kb, 500kb-1Mb, 1-2.5Mb and greater than 2.5Mb) in groups that had roughly the same number of TADs. We found boundaries of TADs in the 500kb-1Mb bin overlapped with gibbon BOS most often, and this was true for regions orthologous to BOS in all other species, except dog **(Supplemental Fig. S9)**. After increasing the resolution of our comparisons by binning TADs into smaller fixed-size bins (incrementing by 200kb), we observed the most significant overlap between BOS and boundaries of TADs in the 400-600kb bin (Figure 4B **and Supplemental Fig. S10)**.

Additionally, we used permutation tests to ascertain whether the TAD boundaries that overlap with BOS are largely the same across species. We found that TADs that overlap with BOS were more conserved than would be expected by chance for the two TAD size groups that we investigated (500kb-1Mb and 400kb-600kb) **(Supplemental Material and Supplemental Fig. S11)**. Focusing on just 500kb-1Mb TADs, we identified 19 loci orthologous to gibbon BOS that overlap with TAD boundaries in all species. Across the non-gibbon species, these loci correspond to 15 regions, since 8 of the 19 BOS derive from reciprocal rearrangements in gibbon and thus map to adjacent locations in non-gibbon species. The 211 genes within +/−500kb of these highly conserved TAD boundaries include genes with essential developmental functions. The mis-regulation of many of these genes has been implicated in lethality in mouse knock-out models and human disorders. Among these is *LYPD6*, whose product is involved in Wnt/ß–catenin signaling and has been found duplicated (microduplication of 2q23.1) or disrupted in autism and other congenital disorders characterized by severe intellectual disabilities (Chung et al., 2012; Nilsson et al., 2017). This gene set also includes *GGN* (gametogenetin), whose complete loss leads to embryonic lethality at the very early stages of pre-implantation due to compromised meiotic double-strand break (DSB) repair (Jamsai et al., 2013). In addition, a recent study reported disruption of three of the highly conserved TADs, which encompasses 36 of the 211 genes in three patients with congenital disorders, including severe developmental impairment (Redin et al., 2017). Furthermore, 161 of the 211 genes (73.6%) are found in a recently curated list of 3,455 disease-associated genes from the Online Mendelian Inheritance of Man (OMIM) (Dickinson et al., 2016) with 34 (21.11%) of them are associated with a disease phenotype. Diseases in this list included retinitis pigmentosa (genes *IFT172*, *ZNF513*, and *RHO*), gastrointestinal defects (*TTC7A*), and Charcot-Marie-Tooth disease, type 1C (*PMP22* and *LITAF*).

### Large new TADs are rarely generated by evolutionary chromosomal rearrangements

Chromosomal rearrangements have the potential to break existing TADs and generate new ones. While this phenomenon has recently been observed in cancer genomes (Dixon, 2017), it has not, to our knowledge, been explored in an evolutionary setting. There are six possible scenarios when the ancestral and current position of rearrangement breakpoints and TAD boundaries are taken into account. In the case in which a BOS co-localize with a TAD boundary in gibbon, the two sites in the ancestral chromosome could have occurred 1) both in boundaries, 2) both inside TADs, or 3) one in a boundary and the other inside a TAD. The same three scenarios can apply when a BOS does not overlap with a TAD boundary in gibbon (Figure 5). When looking at boundaries of TADs sized 500kb-1Mb, we found that 55 out of 67 BOS overlap with TAD boundaries in gibbon; for 31 of these BOS, both sides co-localized with TAD boundaries in the human genome, which represents the ancestral state. This proportion was significantly higher than any of the other scenarios (two-sided McNemar test, p=0.0005) and indicated that evolutionary rearrangements are skewed towards maintaining TADs as intact units. Of the remaining 24 BOS that overlap with TAD boundaries in gibbon, 16 have evolved from breaking DNA at one ancestral TAD boundary and one TAD body. All other 4 evolutionary possibilities have occurred much less often (Figure 5).

**Figure 5.**
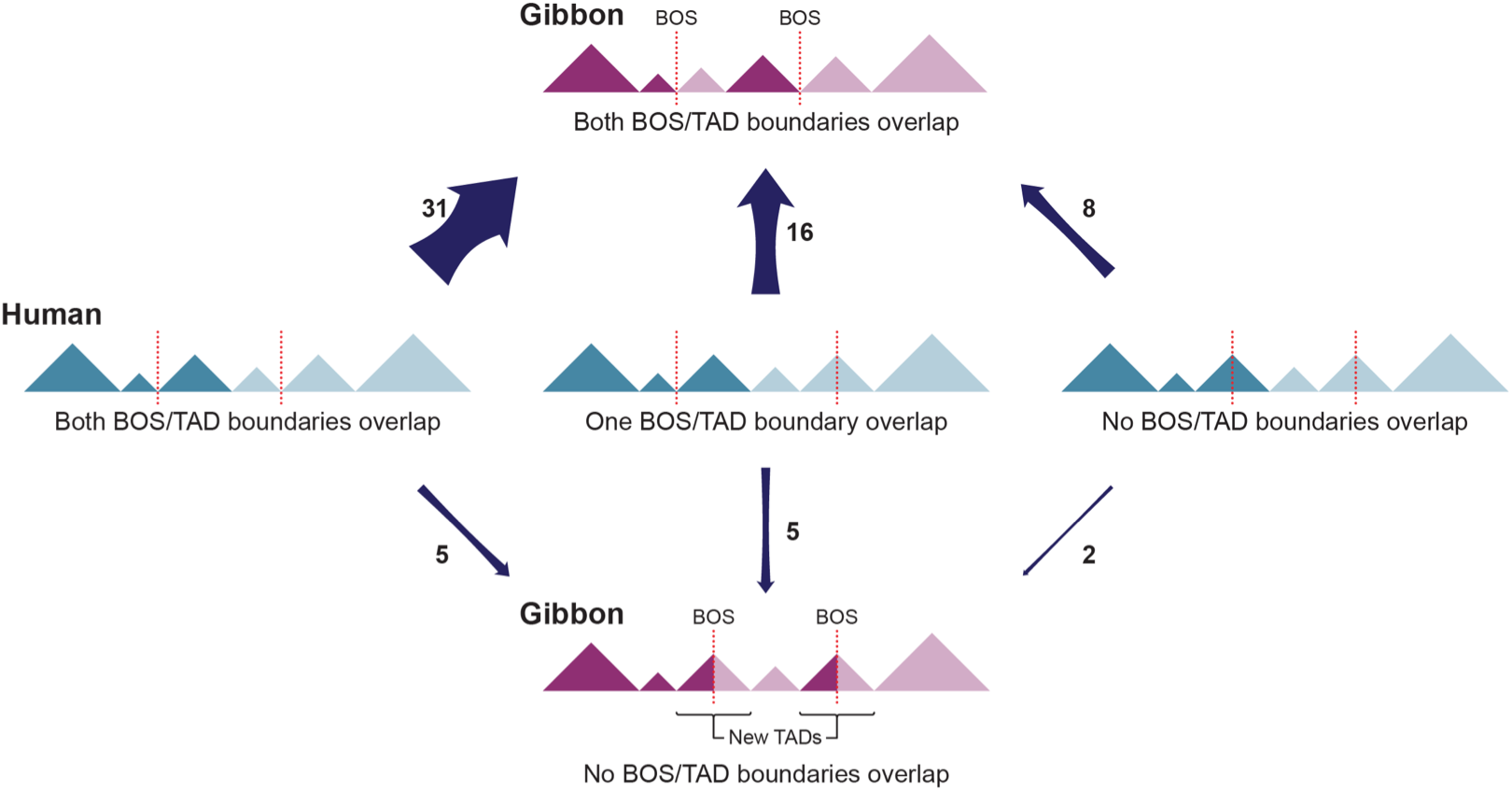
Alternative evolutionary relationship between BOS and TADs. Schematics show all possible scenarios between BOS (dotted lines) and TADs (triangles) in the gibbon (purple) and human (ancestral, blue) genomes. Arrow width relates with prevalence of the scenario in the gibbon genome and the number next to the arrow represents the number of occurrences of each scenario.

Our findings show that it is quite uncommon for gibbon BOS to occur within TAD bodies, away from boundaries. In total, we only found 12 BOS that did not co-localize with TAD boundaries in the gibbon genome (Figure 5). Of these, 7 BOS were located on three gibbon chromosomes (NLE8, NLE11 and NLE13) and 4 of these 7 are results of reciprocal translocations. We found only two gibbon BOS (NLE11_3 and NLE8_3) that did not overlap with boundaries in both gibbon and human. These two rearrangements have broken ancestral TADs and created new ones in the gibbon genome. One of these events involves gibbon chromosomes NLE10 and NLE11, which originated from a reciprocal translocation between ancestral chromosomes corresponding to human 12 and 19 (Figure 6A). On human chromosome 19, this rearrangement broke a cluster of highly transcribed Kruppel-associated box (KRAB)-containing Zinc-finger-containing (KZFP) genes (Dai et al., 2003) bringing a portion of the ZNF cluster together with *TBCID30*, *GNS* and *RASSF3* within a gibbon TAD (Figure 6A). Because the breakpoint on NLE11 does not overlap with a boundary, this rearrangement could cause the regulatory landscape of the ZNF cluster to merge with the one of the downstream genes (*GNS* and *RASSF3*).

**Figure 6.**
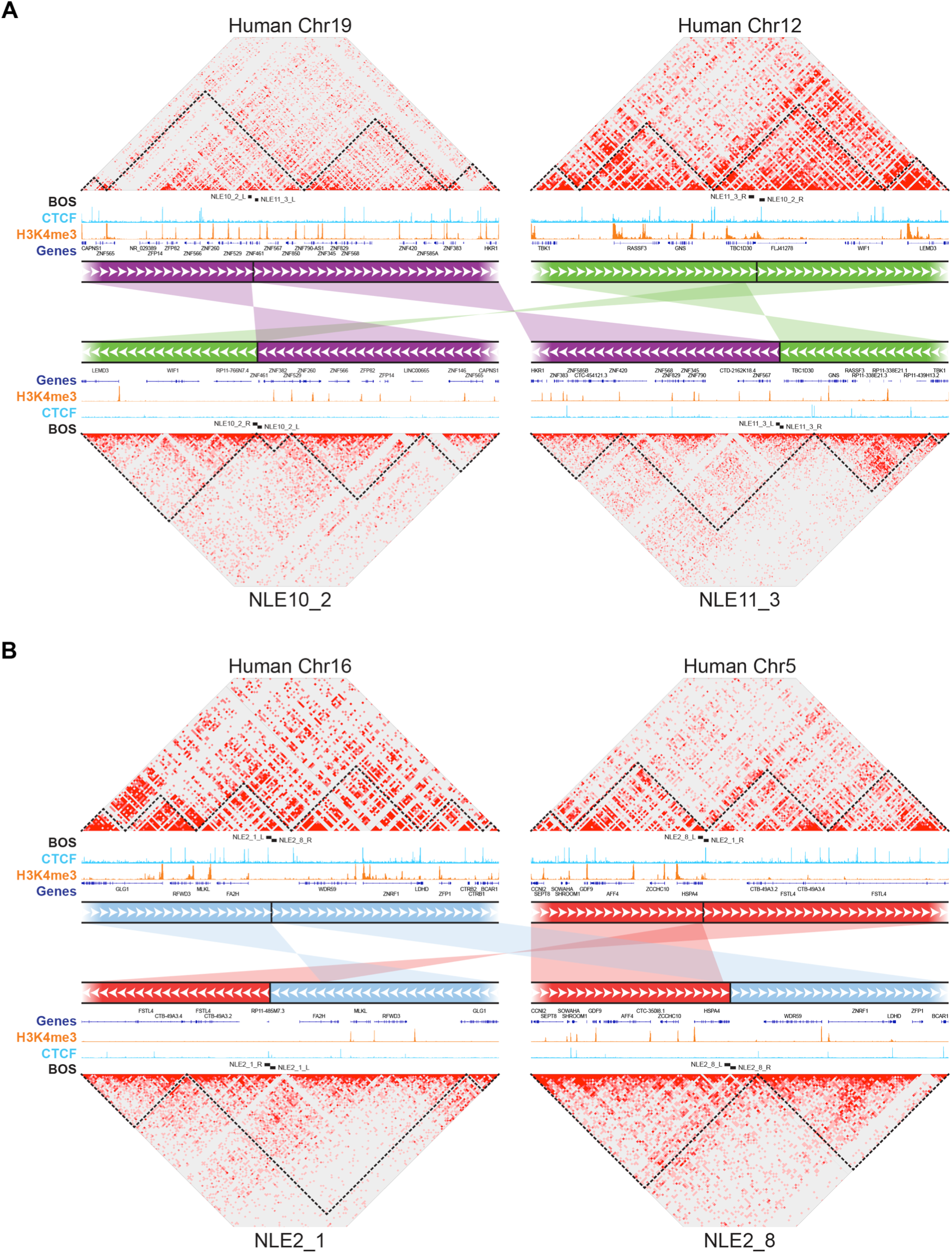
New TADs can originate from genomic rearrangements. A) Example of a reciprocal translocation and inversion whose breakpoints (NLE10_2 and NLE11_3) do not overlap with TAD boundaries in human (top). NLE11_3 (bottom left) maps within a TAD in gibbon (bottom), but it is close to an ancestral boundary, while NLE10_2 (bottom right) corresponds to a new boundary on NLE10. ChIP-seq pileups for H3K4me3 (orange) and CTCF (light blue) are shown for human (top, ENCODE) and gibbon (bottom). B) Example of reciprocal translocations in which BOS (NLE2_1 and NLE2_8) are both in TAD boundaries in human (top) and but not in gibbon (bottom). Although not overlapping, NLE2_8 is located very close to TAD boundaries in gibbon while a new TAD was created by the rearrangement involving NLE2_1.

However, this effect may have been mitigated by an ancestral TAD boundary marked by a CTCF binding event upstream the BOS (Figure 6A). Looking at the reciprocal rearrangement formed by this event, our analysis revealed that the breakpoint on NLE10 (NLE10_2) corresponded to a new TAD boundary (i.e. present in gibbon, but not in human or rhesus) that insulates the active ZNF cluster from other genes deriving from human chromosome 12 (*WIF1* and *LEMD3*). In five of the 67 BOS we found, breakpoints that occurred at ancestral (human) TAD boundaries but that are not boundaries in the gibbon genome. By generating new gibbon-specific TADs these types of rearrangements might lead to functional novelty. When inspecting the Hi-C data, we found that three of these BOS (NLE2_8, NLE20_1, and NLE11_4) have a TAD boundary located nearby. One of these cases is the reciprocal translocation between chromosomes corresponding to human 16 and 5, followed by an inversion that gave rise to NLE2 (Figure 6B). The breakpoints of this inversion (NLE2_1 and NLE2_8) do not perfectly co-localize with TAD boundaries in gibbon, but NLE2_8 is very close to the next boundary (~30kb) with no genes present in between. In the other two instances (NLE14_13 and NLE2_1), a new gibbon TAD has been created. Overall, these observations indicate a lack of new TADs in gibbon and that emergence of new TAD boundaries to prevent ectopic interactions and gene mis-regulation when TAD structures are disrupted (Dixon, 2017; Lupianez et al., 2016).

### The chromatin on the two sides of gibbon BOS maintain their ancestral epigenetic identity

Few studies have explored the extent of epigenetic remodeling that occurs after evolutionary chromosomal rearrangements. We asked whether the two sides of gibbon BOS display different epigenetic states, as may be expected from distal non-related chromatin, or have undergone epigenetic homogenization after becoming contiguous on the rearranged chromosome. To characterize the epigenetic landscape of gibbon BOS, we generated whole-genome bisulfite sequencing (WGBS) data from whole blood of Vok **(Supplemental Material)**. To normalize for effects of CpG density, we calculated residual methylation after fitting logistic functions to methylation, given CpG density **(Supplemental Fig. S4)**. We observed large differences in the distribution of CpG methylation, CpG density, and H3K4me3 peaks between the two sides of BOS, often with a sharp switch occurring at the breakpoint (Figure 7A).

**Figure 7.**
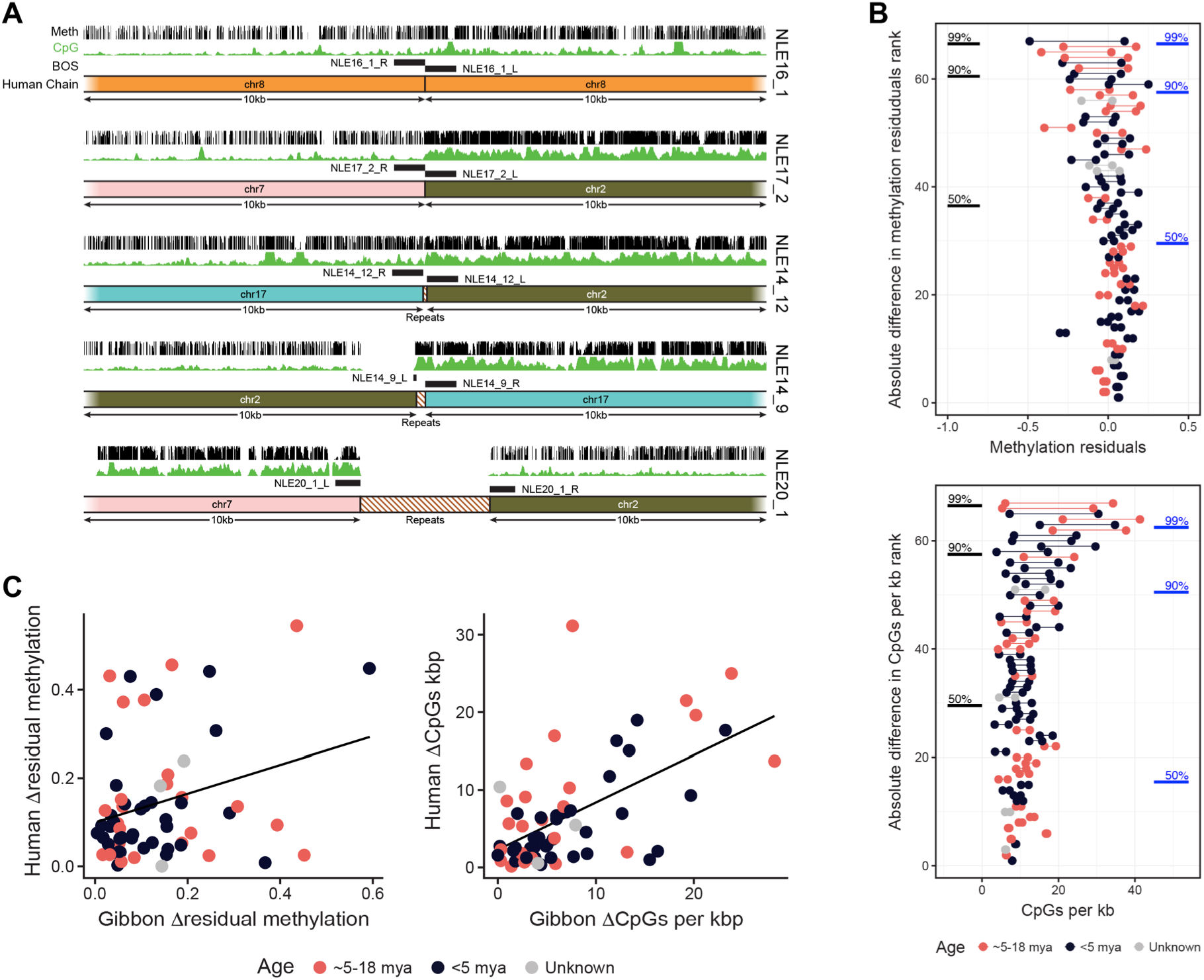
Gibbon BOS maintain their ancestral epigenetic identity and resemble non-syntenic regions. A) Examples of BOS showing a noticeable difference in CpG density (green track) and methylation (black track) between the two sides of the rearrangement with the switch occurring at the BOS (black blocks). Homology with the human chromosomes is shown below each BOS. Gibbon-specific repeats within the breakpoint explain the gap with the human alignment. (NLE = *Nomascus leucogenys*; Meth = methylation). B) Ranked barbell plots show the difference in residual methylation and CpG density between the two sides of each of the gibbon BOS. Each point represents a BOS side and a line segment joins the two sides from the same BOS. BOS are ordered vertically by magnitude of the difference between sides. Black lines on the left show the rank associated with percentiles of distal permutation regions, while blue lines on the right show ranks for percentiles for adjacent permutation regions. Color-coding by age of the rearrangement highlights that old BOS (5-18 mya) are as likely as young ones (<5 mya) to show a large difference between the two sides. C) Scatter plots of Δ residual methylation (left) and Δ CpG density (right) between gibbon and human BOS regions where each point represents one BOS. The line shows a least squared linear regression and the points are color-colored as in B.

To quantify this observation, we calculated the absolute difference in mean residual CpG methylation and CpG density (Δmeth and ΔCpG) between the two sides of each BOS (+/−10kb) and used permutation analyses to evaluate whether the Δ values observed in BOS were greater than would be expected by chance. We performed the same analysis for H3K4me3 peaks (ΔH3K4me3), but with a larger window around BOS (+/−500kb) to account for the lower density of the peaks **(Supplemental Material and Supplemental Fig. S12)**. We found that despite being contiguous on gibbon chromosomes, the two sides of BOS are significantly different in their epigenetic characteristics and behave like regions distantly located in the gibbon genome **(Supplemental Table S2)**. Leveraging WGBS data already available for human (Hernando-Herraez et al., 2015), and generated by us for rhesus, we used regions orthologous to the gibbon BOS to reconstruct *in silico* the arrangements found in gibbon and repeated the permutation analysis in these species. As expected, in both human and rhesus CpG density and DNA methylation between the two sides of BOS look as different as distal regions **(Supplemental Table S2)**. To test whether the epigenetic landscape across BOS homogenizes with time, we split gibbon BOS based on their cytogenetically determined age (Capozzi et al., 2012) into ancestral (5-18 mya old) or Nomascus-specific (<5 mya old) when possible. We found no clear relationship between age of BOS and difference in residual methylation and CpG density across BOS, as older BOS also show large Δmeth and ΔCpG (Figure 7B).

Very similar results were obtained for human and rhesus **(Supplemental Fig. S13A)** and the amount of epigenetic difference measured for each BOS is highly correlated between species (Pearson correlations for gibbon vs. human ΔCpG: 0.57, Δmeth: 0.31; gibbon vs. rhesus ΔCpG: 0.70, Δmeth: 0.44), suggesting that the epigenetic differences across BOS in inherited from the primate common ancestor (Figure 7C **and Supplemental Fig. S13B)**. Overall, these results indicate that chromosomal regions largely maintain their ancestral epigenetic landscape after rearrangement, and that physical contiguity at BOS does not lead to epigenetic homogenization over time.

## Discussion

Recent studies have demonstrate how structural variations in the genome can alter DNA interaction and topologically associating domains (TADs) in the context of pathology and disease (Franke et al., 2016; Hnisz et al., 2016; Lupianez et al., 2015; Nilsson et al., 2017; Sun et al., 2017). However, the consequences of chromosomal rearrangements during genome evolution have not been extensively explored. In this study, we use a high-resolution map of gibbon-human breaks of synteny (BOS) from the heavily reshuffled gibbon genome and genetic and epigenetic data to thoroughly examine the evolutionary relationship between chromosomal rearrangements and structure of TADs.

Using gibbon Hi-C data, we visualized genome-wide DNA interactions and detected multiple non-reference rearrangements, the most evident of which was experimentally validated and represented a known polymorphic translocation (Figure 2A) (Koehler et al., 1995). We also identified unusual long-distance “ghost interactions”, resulting from a very recent (~2 mya) pericentromeric inversion in NLE7 (Figure 2C). Ghost interactions, which to our knowledge have not been described before, indicate that preservation of long-distance DNA interactions in the 3D space may be a transitional solution to maintain the regulatory networks after a large-scale genomic rearrangement has occurred. It is not clear how these ghost interactions are maintained and for how long, but further analysis of recent or polymorphic rearrangements in the human population and other very recent evolutionary rearrangements, may help shed some light on this phenomenon.

We used our Hi-C interaction data to computationally predict TADs in the gibbon genome and compare their location relative to regions of gibbon-human BOS. We found that BOS consistently and significantly co-localize with TAD boundaries in the gibbon genome and that this association was present independent of the parameters used to predict gibbon TADs **(Supplemental Fig. S8)**. Furthermore, gibbon BOS showed significant enrichment of genetic and epigenetic signatures of TAD boundaries, including higher CpG density than the rest of the genome, enrichment in CTCF binding, H3K4me3, and presence of SINE elements (Figure 3B **and Supplemental Fig. S6)**. In most cases, the two sides of the gibbon BOS corresponded to TAD boundaries in the inferred ancestral hominoid state (Figure 5) demonstrating that most TADs were maintained as intact modules during and after rearrangement. This overlap was, however, observed more often for larger TADs (500kb-1Mb) suggesting that smaller sub-TADs are more variable during evolution (Figure 4B **and Supplemental Fig. S9)**. Leveraging publicly available data, we also showed that large TADs that overlap with BOS are often conserved across gibbon, human, rhesus macaque, mouse, dog and rabbit (Figure 4B **and Supplemental Fig. S10)** and therefore they pre-existed and survived genome remodeling in gibbon.

Two, non-mutually exclusive, models can explain the striking co-localization of BOS and TAD boundaries. The “fragile TAD boundary” model predicts that the distinct genetic and epigenetic properties of TAD boundaries along with their open and transcriptionally active chromatin state, elicit a higher rate of DNA double strand break (DSB) and repair (Figure 8) (Smerdon, 1991). In support of this model, open chromatin and the epigenetic marks found at TAD boundaries (e.g. H3K4me3 and CTCF) have been linked to chromatin fragility in human disease (De and Michor, 2011; Li et al., 2012; Tchurikov et al., 2015), species evolution (Carbone et al., 2009a; Lemaitre et al., 2009) and simulation analyses (Berthelot et al., 2015). Moreover, an *in vitro* study recently showed that chromosome loop anchors bound by CTCF and cohesin are vulnerable to DSBs mediated by topoisomerase 2B (TOP2B) and act as fragile sites for chromosomal rearrangements (Canela et al., 2017). Thus, higher frequency of breakage and repair at TAD boundaries relative to other genomic regions could explain the skewed distribution of rearrangements in gibbon and other species. Under this model, we would also expect to observe a similar trend in somatic (e.g. cancer cell lines) and germline rearrangements in the human population. Indeed, it was recently discovered that nearly all short tandem repeats (STRs) linked to repeat expansion diseases co-localize with TAD boundaries and that their expansion severely compromises those boundaries in Fragile X Syndrome and Huntington’s disease (Sun et al., 2017).

**Figure 8.**
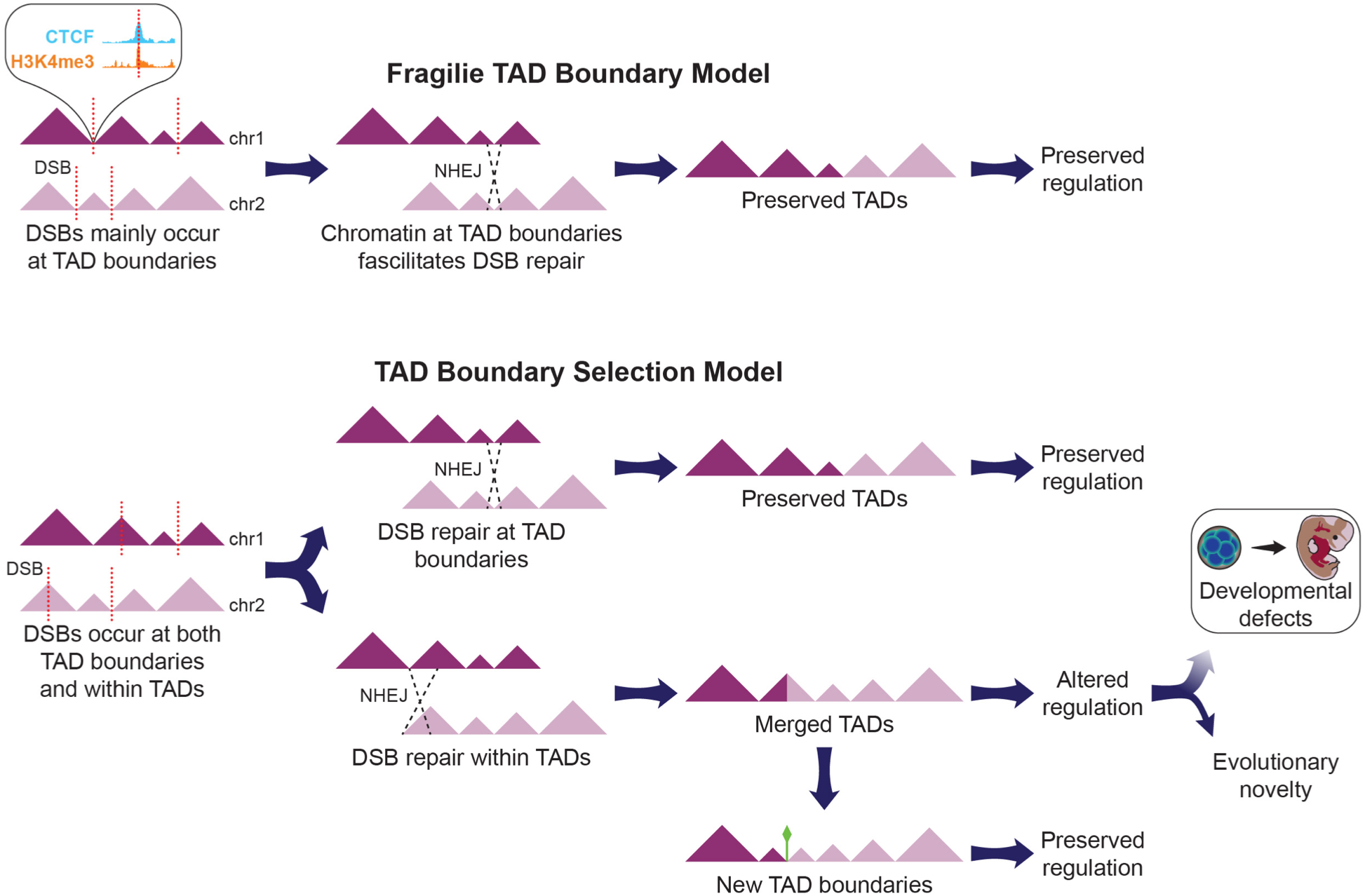
Two models to explain co-localization of BOS and TAD boundaries in genome evolution. A) Based on the “fragile TAD boundary” model, TAD boundaries carry epigenetic marks associated with DNA double strand break (DSB, red dotted lines) and repair. DSBs will therefore occur more frequently at TAD boundaries than other genomic regions and have a higher chance to be repaired and evolutionarily fixed. B) The “TAD boundary selection” model assumes that DSBs occur equally at TAD boundaries and within TADs. However, rearrangements altering TAD structure by misplacing or deleting TAD boundaries are lost through purifying selection, while those maintaining TADs intact are more likely to become evolutionary fixed. In a small portion of the cases, new TAD boundaries might emerge (green diamond) and survive in the population.

In the “TAD boundary selection” model, DSBs are predicted to occur equally at TAD borders and within TAD bodies. However, DSBs that co-localize with boundaries and maintain boundary function are more likely to become evolutionarily fixed, while DSBs elsewhere likely reduce fitness and are eliminated by purifying selection (Figure 8). In agreement with this model, we showed that TAD boundaries in correspondence of BOS have maintained their insulating function and prevented epigenetic homogenization between the two sides of the rearrangements in the gibbon genome (Figure 7A-B). We also revealed that highly conserved boundaries were located near important developmental genes, suggesting that rearrangements that cause mis-regulation of these genes were likely negatively selected. Our findings suggest that both models may have been at play during the evolution of the gibbon genome, but further studies are required to test these paradigms, including directly examining TAD boundary fragility *in vitro* (Canela et al., 2017) andtesting selection models across different species with rearranged genomes.

While the majority of our gibbon BOS co-localized with TAD boundaries, a small subset of the rearrangements created new TADs that were evolutionarily fixed. Many of these new gibbon TADs appear to mitigate ectopic regulation by establishing new boundaries to prevent interactions between the newly joint genomic regions, especially when they present very different regulatory landscapes. For example, on gibbon chromosome 10, genes *WIF1* and *LEMD3* were joined by a reciprocal translocation to a transcriptionally active cluster of zinc finger genes and a new TAD boundary was formed in gibbon nearby the BOS (Figure 6A). This new boundary might prevent ectopic activation of *LEMD3* whose mutations have been associated with skeletal dysplasia and collagen-type nevi (Bushke-Ollendorff syndrome) and rare mesenchymal dysplasia (Melorheostosis) (Debeer et al., 2003; Fischetto et al., 2017; Hellemans et al., 2004). Alternatively, new TADs might become evolutionarily fixed because they are advantageous. For instance, domain reorganization may bring distant, co-regulated genes into the same TAD and facilitate their coordinated expression (Harewood and Fraser, 2014; Schoenfelder et al., 2010). We made some anecdotal observations in support of this scenario. On gibbon chromosome 8 a new TAD originated from a translocation that joined a potassium channel gene (*KCNH8)* and a vacuolar ATPase (*ATP6V1B2*), both highly expressed in the human brain (Consortium, 2013). Mutations in *ATP6V1B2* cause a developmental disorder known as Zimmermann-Laband syndrome (ZLS); the same phenotype is also observed when mutations occur independently in the gene *KCNH1* (Kortum et al., 2015). The overlap in clinical features for these two physically distant genes with apparently unrelated functions was explained by the fact that the two proteins have coordinated action: KCNH1 provides K+ as a counter-ion that is needed by *ATP6V1B2* to pump protons and achieve acidification of intracellular vacuoles (Kortum et al., 2015). We therefore speculate that similar coordinated functions exist between *KCNH8* (whose function and expression are very similar to *KCNH1)* and *ATP6V1B2* and that co-localization of the two genes in the new TAD in gibbon has facilitated their co-expression. Similarly, we observed joining of two genes of the RAS oncogene GTPases (*RAB37* and *RAB40B*) in a new TAD on NLE14, and two genes (*HMX1* and *ADRA2C*) implicated in regulating the function of sympathetic neurons (Furlan et al., 2013) in a new TAD on NLE20 (data not shown). Overall, these preliminary observations suggest that new TADs that either mitigate ectopic interaction or facilitate co-expression of genes with related function are more likely to be tolerated and fixed in the genome.

To determine if any epigenetic remodeling and homogenization occurs in the genome after chromosomal rearrangements, we compared aspects of the epigenetic landscape across the gibbon BOS. We measured both residual DNA methylation and H3K4me3 distribution, as indicators of repressed and active chromatin respectively, and found large differences in chromatin state between the two sides of the rearrangements with the transition occurring exactly at the BOS (Figure 7C). The overall epigenetic state of chromatin around gibbon BOS was strongly correlated to orthologous regions in human and rhesus **(Supplemental Fig. S13)**, thus the original epigenetic state remained mostly stable, even in the oldest gibbon BOS (Figure 7B). Future studies should compare transcription levels and other functional epigenetic marks (e.g. H3K27ac, which marks enhancers), to examine local regulatory and transcriptional changes that might occur even when the overall chromatin state remains unchanged. Nevertheless, our findings demonstrate interesting insight into the preservation of epigenetic landscapes despite extensive evolutionary genomic rearrangements.

In conclusion, this study is the first to show a remarkable correspondence between evolutionary breakpoints and TAD boundaries in the gibbon genome, a critically endangered primate species that recently experienced rapid and heavy chromosome remodeling (Carbone et al., 2014). Our findings provide compelling evidence that this co-localization may be due to both TAD boundaries’ higher fragility and easier repair, as well as purifying selection against rearrangements that disrupt regulation of essential TADs (Figure 8). Formation of new large-scale TADs does not appear to play a large role in emergence of evolutionary novelty in the gibbon genome. New TADs seem to be more tolerated when ectopic interactions are mitigated through formation of new boundaries, or alternatively, when genes with coordinated expression are placed in the same TAD. Finally, we find that the two sides of evolutionary breakpoints remain epigenetically similar to their ancestral state and do not homogenize even after millions of years. Overall, this study supports a non-random mode of chromosome evolution, where functional genomic units remain genetically and epigenetically intact, despite being shuffled around in the genome.

## Methods

### Annotation of gibbon BOS and orthologous BOS in other species

We compiled a comprehensive list of gibbon-human BOS by incorporating all previous findings from array painting (Carbone et al., 2006), fluorescent *in situ* hybridization (FISH) experiments, end-sequencing of Bacterial Artificial Chromosomes (BACs) (Girirajan et al., 2009; Roberto et al., 2007), shotgun sequencing, assembly of full BACs (Carbone et al., 2009a; Carbone et al., 2006), and the latest gibbon genome assembly, Nleu3.0 (Carbone et al., 2014). We created a naming convention to account for the directionality of the rearrangement and distinguish between the two sides (*e.g*. NLE5_2_R_r, indicates the right side (R) of the second BOS on gibbon chromosome 5, extended 10kb downstream (r)). To enable comparative analyses, we identified regions orthologous to gibbon BOS in human, rhesus macaque, mouse, dog, and rabbit by using BLAT and the UCSC genome browser LiftOver tool (Kent et al., 2002). From the 67 gibbon BOS, we identified 133 orthologous regions mapped on human (*Homo sapiens*, hg38), 129 on rhesus (*Macaca mulatta*, RheMac8), 95 on mouse (*Mus musculus*, GRC38), 104 on rabbit (*Oryctolagus cuniculus*, oryCun2), and 114 in dog (*Canis familiaris*, CanFam3.1) **(Supplemental Table 1)**.

### Genome-wide chromatin conformation capture (Hi-C) sequencing and analysis

To prepare the Hi-C library, 2.5 x 10^7^ *Vok* EBV-transformed cells (Supplemental Material) were cross-linked with 1% formaldehyde for 5 minutes on ice and then lysed in Lysis Buffer (0.1% SDS, 0.5% Triton X-100, 20mM Tris-HCl pH 8.0, 150mM NaCl and protease inhibitor cocktail (Roche)). The chromatin was pelleted and washed twice with Hi-C Wash Buffer (HWB; 50mM Tris-HCl pH 8.0, 50mM NaCl), then re-suspended in 250μl HWB buffer with 0.6% SDS at 68°C for 10 minutes. Replicates of 50μL lysate (<2μg chromatin per replicate) were bound to 100 μL AMPure XP beads (Beckman Coulter), then washed twice with HWB. Bead-bound chromatin was digested in NEB DpnII Buffer with 5U of DpnII enzyme for 3 hours while shaking at 37°C. After washing beads with HWB, biotinylated dCTP was incorporated by incubating beads in 50μL of End Fill Mix (NEB Buffer 2, 0.15mM each dATP, dGTP, dTTP (New England Biolabs), 0.04mM biotin-14-dCTP (Invitrogen), 3.75U Klenow (New England Biolabs) for 30 min while shaking at 25°C. Beads were washed with HWB and then chromatin proximity ligation was carried out by overnight incubation at 16°C in a 250μL reaction containing T4 DNA Ligase Buffer (New England Biolabs), 0.1mg/ml BSA (New England Biolabs), 0.25% Triton X-100 (Fisher), and 50U T4 DNA Ligase (New England Biolabs). Next, 2.5μL 10mM dNTPs (New England Biolabs) and 7.5U T4 DNA Polymerase (New England Biolabs) were added to remove biotin-dCTP from unligated ends.

The samples were released from the beads in 50μL cross-link reversal buffer (50mM Tris pH=8.0, 1% SDS, 0.25mM CaCl_2_, and 0.5 mg/mL Proteinase K) by subsequent incubations at 55°C then 68°C. After cross-link reversal, the samples were cleaned with a 2X AMPure XP bead clean up. DNA was quantified by Qubit fluorometer before Illumina sequencing library preparation using the NEBNext Ultra II DNA Library Preparation Kit (New England Biolabs), following manufacturer’s protocol. The ligation product was pulled down using 25 μL Dynabeads MyOne Streptavidin C1 (Invitrogen) that had been washed with TWB (10mM Tris-HCl pH 8.0, 0.5mM EDTA, 0.05% Tween-20) and suspended in NTB (10mM Tris-HCl pH 8.0, 2M NaCl, 1mM EDTA). The biotinylated ligation junctions were captured with the prepped Dynabeads and then washed with LWB (10mM Tris-HCl pH 8.0, 1M LiCl, 1mM EDTA, 0.05% Tween-20), NWB (10mM Tris-HCl pH 8.0, 0.1M NaCl, 1mM EDTA, 0.05% Tween-20) and HWB before continuing on to indexing PCR. Libraries were multiplexed and sequenced on the Illumina NextSeq 500 platform at the OHSU Massive Parallel Sequencing Shared Resources (MPSSR) to generate 75bp paired-end reads.

Hi-C datasets for the other species: rhesus, mouse, rabbit, and dog and human were obtained from public sources (Grubert et al., 2015; Vietri Rudan et al., 2015). When the public sources contained multiple read files and replicates, we combined data across replicates. All Hi-C datasets were processed similarly using the Juicer pipeline (Durand et al., 2016). For each species, we used TADtool (Kruse et al., 2016) to call TADs using the directionality index method (Dixon et al., 2012) with 140 parameter combinations: seven resolutions (10kb, 25kb, 50kb, 100kb, 250kb, 500kb and 1Mb), five significance cutoffs (0.1, 0.05, 0.01, 0.005 and 0.001) and four window sizes (1Mb, 2Mb, 5Mb and 10Mb). Finally, we used Homer v4.9 (Heinz et al., 2010) to visualize the Hi-C contact maps overlapped across BOS regions.

### Chromatin immunoprecipitation sequencing (ChIP-seq) and data analysis

CTCF ChIP-seq data generation and analysis have been described previously (Carbone et al. 2014). To generate H3K4me3 ChIP-seq libraries, we fixed 5 x 10^7^ cells from EBV-transformed line generated for *Vok* and *Asia* (the individual used for the gibbon genome assembly) with 1% formaldehyde on ice for 5 minutes. Next, cells were lysed with Lysis Buffer (0.1% SDS, 0.5% Triton X-100, 20mM Tris-HCl pH 8.0, 150mM NaCl and protease inhibitor cocktail (Roche)). The lysate was sheared using the Bioruptor Plus sonicator for one cycle of 15 minutes (30 seconds on/off) at high power. A 10ul aliquot of lysate was taken as ”input”. Lysate was split into two preps and each sample was incubated with 1μg of anti-H3K4me3 antibody (abcam) overnight at 4°C. Prepared Pierce Protein A/G Magnetic Beads (ThermoFisher) were added to lysates, incubated for 2 hour at 4°C and then washed four times with TBST (25mM Tris-HCl pH 7.6, 250mM NaCl, 0.05% Tween-20), once with LiCl Wash Buffer (10mM Tris-HCl pH 8.0, 0.5M LiCl, 1mM EDTA, 0.05% Tween-20), once with Lysis Buffer and once with 1X TE. Following washes, samples were eluted with Elution Buffer (1% SDS, 0.1M NaHCO3) and incubated at room temperature for 15 minutes. Elution reactions were pooled and incubated with NaCl overnight at 65°C to reverse crosslinks. Following RNase A and Proteinase K digests at 37°C and 55°C, respectively, samples were cleaned using standard Phenol:Chloroform cleanup and a 1.6X AMPure XP (Beckman-Coulter) bead clean up. 10ng of ChIP and input DNA were used to make next-generation sequencing libraries using the NEBNext Ultra II DNA Library Preparation Kit (New England Biolabs). Libraries were checked with Qubit fluorometer and Bioanalyzer (Agilent) assays and sequenced on the Illumina NextSeq 500 platform to generate single-end 75bp reads.

H3K4me3 ChIP and input sequences (single-end 36bp reads) for rhesus macaque were obtained from public data (Accession number GSE60269) (Zhao et al., 2006). For both gibbon and rhesus data, we removed adapter sequences using trimmomatic v0.35 (Bolger et al., 2014) and de-duplicated reads using fastx-toolkit v0.0.13 (Hannon). We then mapped the reads to their corresponding genomes with BWA-mem v0.7.9a (Li and Durbin, 2009) and removed reads with mapping scores below 30. Next, we used phantompeakqualtools v2.0 (Marinov et al., 2014) to run SPP (Kharchenko et al., 2008) and obtain cross-correlation profiles and call a loose set of 300,000 peaks. In order to determine a cutoff for these peaks, we performed an irreproducibility discovery rate (IDR) analysis (Landt et al., 2012). Using an IDR cutoff of 0.0025 (recommended by ENCODE (Consortium, 2012)) we obtained 20,828 peaks for gibbon and 21,738 peaks for rhesus.

### Whole genome bisulfite sequencing (WGBS) and data analysis

Whole genome bisulfite-converted samples were generated for gibbon and rhesus as previously described (Lister et al., 2009). Briefly, 1μg genomic DNA was extracted from opportunistic whole blood samples from *Vok* and rhesus and fragmented (Fragmentase Reaction Buffer v2, Fragmentase (New England Biolab)), end-repaired (T4 DNA Ligase Buffer, dNTP, T4 DNA polymerase, T4 PNK), A-tailed (NEB buffer 2, 1mM dATP, Klenow exo-) and ligated to 30μM methylated forked Illumina adaptors (Quick Ligase Reaction Buffer, Quick Ligase). Ligated fragments were size-selected (250-500bp) and bisulfite converted using the Qiagen EpiTect bisulfite kit. Fragments were amplified and sequenced on the Illumina HiSeq2000 platform to generate 111,876,578 paired-end reads. Human WGBS data was publicly available (Accession SRR2058107 and SRR2058108) (Hernando-Herraez et al., 2015).

Raw reads from human, rhesus, and gibbon were pre-processed using trim_galore v0.4.0 (Krueger and Andrews, 2011) to remove adapters, bases with Phred score <20 and reads shorter than 20bp. Next, we used bsmooth v0.8.1 (Hansen et al., 2012) to map 250,000 random reads from each dataset to their corresponding genomes and generate mbias plots to determine whether the methylation level correlates with the position of the CpG in the reads. Based on these plots and the output of FASTQC v0.10.1 (Andrews) quality reports, we uniformly trimmed 5bp from both ends of all reads. We then mapped reads to their respective genomes using Bismark v0.12.5 (Krueger and Andrews, 2011) and obtained methylation calls for each read at each CpG locations. In the subsequent analysis, the counts of all CpGs were assessed, but methylation values were only considered for CpG covered by >4 reads. For the human sample, data from two sequencing lanes was merged at the CpG summary level.

### Data Access

All scripts are available at github.com/nathanlazar/APE_METH_bin Data generated for this project are pending SRA submission and are currently available under request.

## Acknowledgments

We are thankful to the Gibbon Conservation Center (GCC) for providing us with the samples used in this study. In particular, we thank Gabriella Skollar for her inspiring work and passion taking care of these critically endangered species. Illumina sequencing was performed by the OHSU Massively Parallel Sequencing Shared Resource (MPSSR). We also thank the OHSU ExaCloud Cluster Computational Resource and the Advanced Computing Center that allowed us to perform the intensive large-scale data workflows. We thank Dr. Suzi Fei and Melissa Yang for helping deposit the sequences, Brett Davis for helping with mapping WGBS data, and Dr. Rachel O’Neill for performing the FISH experiment. Finally, we thank Drs. Andrew Adey and Eric Jorgenson for insightful discussions about this manuscript. N.H.L. and T.J.M. were supported by a fellowship from the National Library of Medicine Biomedical Informatics Research Training Program T15 LM007088during this project; L.C. is supported by NIH/NCRR P51 OD011092, NSF 1613856, and the Knight Cardiovascular Institute.

## Author Contributions

N.H.L. and L.C. designed the experiments and the analysis plan. N.H.L. performed the analysis. K.A.N., L.C performed the experiments. B.O. and R.E.G. contributed to experimental methods. L.C., N.H.L., and M.O. wrote the paper. T.J.M. created the figures. All authors were involved with editing the manuscript.

